# A synthetic Calvin cycle enables autotrophic growth in yeast

**DOI:** 10.1101/862599

**Authors:** Thomas Gassler, Michael Sauer, Brigitte Gasser, Diethard Mattanovich, Matthias G. Steiger

## Abstract

The methylotrophic yeast *Pichia pastoris* is frequently used for heterologous protein production and it assimilates methanol efficiently via the xylulose-5-phosphate pathway. This pathway is entirely localized in the peroxisomes and has striking similarities to the Calvin-Benson-Bassham (CBB) cycle, which is used by a plethora of organisms like plants to assimilate CO_2_ and is likewise compartmentalized in chloroplasts. By metabolic engineering the methanol assimilation pathway of *P. pastoris* was re-wired to a CO_2_ fixation pathway resembling the CBB cycle. This new yeast strain efficiently assimilates CO_2_ into biomass and utilizes it as its sole carbon source, which changes the lifestyle from heterotrophic to autotrophic.

In total eight genes, including genes encoding for RuBisCO and phosphoribulokinase, were integrated into the genome of *P. pastoris*, while three endogenous genes were deleted to block methanol assimilation. The enzymes necessary for the synthetic CBB cycle were targeted to the peroxisome. Methanol oxidation, which yields NADH, is employed for energy generation defining the lifestyle as chemoorganoautotrophic. This work demonstrates that the lifestyle of an organism can be changed from chemoorganoheterotrophic to chemoorganoautotrophic by metabolic engineering. The resulting strain can grow exponentially and perform multiple cell doublings on CO_2_ as sole carbon source with a µ_max_ of 0.008 h^−1^.

**Graphical Abstract:** 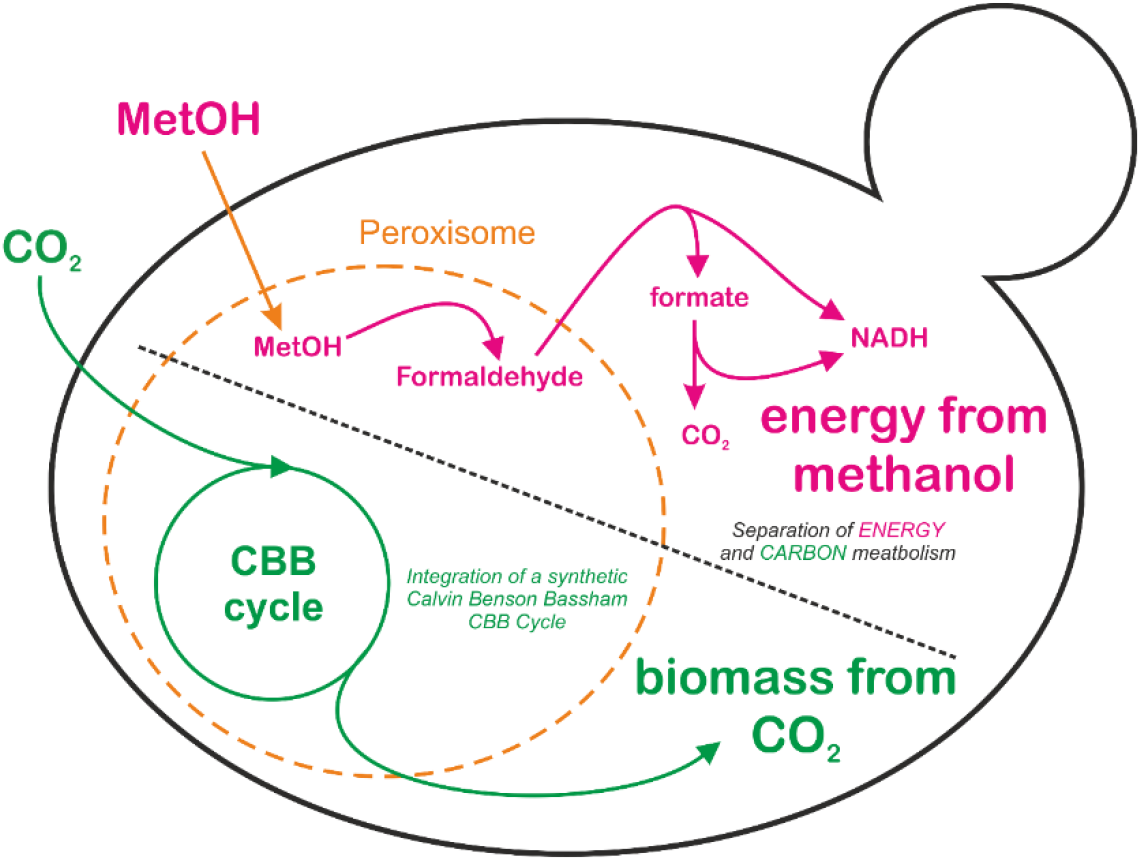

## 1. Introduction

Autotrophic organisms have the capability to use inorganic CO_2_ as a carbon source to build up their biomass. Inorganic carbon assimilation is essential for a closed carbon cycle on earth and thus supplying heterotrophic organisms with organic carbon sources. The excessive use of fossil resources by mankind led to a global imbalance of the carbon cycle leading to increasing atmospheric CO_2_ concentrations and to global warming. Finding new ways to reduce atmospheric CO_2_ requires a better understanding and utilization of the metabolic pathways for assimilation of CO_2_ into biomass. Biotechnological production hosts are almost exclusively heterotrophs and use feedstocks which compete with feed or food production. One of the key tasks for creating sustainable biotechnological processes in the near future is to use CO_2_ as a carbon feedstock and thereby reducing the atmospheric levels of this greenhouse gas. The Calvin-Benson-Bassham (CBB) cycle ^1^ is the predominant of the six naturally occurring CO_2_ fixation pathways^2,3^. Its key enzyme ribulose-1,5-bisphosphate carboxylase/oxygenase (RuBisCO) is considered to be the most abundant enzyme found in the biosphere and fixates around 90% of the inorganic carbon converted into biomass ^4^.

Engineering efforts were made to equip heterotrophic model organisms with CBB cycle genes including *Escherichia coli* and *Saccharomyces cerevisiae*. Expressing RuBisCO and phosphoribulokinase (Prk) in *E. coli* led to a decreased exhaust of CO_2_ when growing on arabinose ^5^. An important step was achieved by splitting the carbon metabolism of *E. coli* into modular subunits: one providing energy and reducing equivalents in the form of ATP and NADH, and the other module carrying out the carbon assimilation. As an energy source Antonovsky and coworkers used pyruvate to fuel a synthetic CBB cycle in *E. coli* enabling “hemiautotrophic” growth ^6^. The term hemiautotrophic was introduced because only part of the carbon assimilated originated from CO_2_ and the rest came from pyruvate. In a follow-up study it was shown which mutations *E. coli* requires to stabilize the metabolic network in order to host a non-native metabolic pathway as the CBB cycle ^7^. Implementing parts of the CBB cycle in the yeast species *S. cerevisiae* ^8–11^ increased yields in ethanol production by using CO_2_ as an additional electron acceptor to reoxidize NADH. In a recent study, the assimilatory pathway of *Methylobacterium extorquens AM1* was blocked by deleting three native genes (*ccrΔ*, *glyAΔ* and *ftfLΔ*). RuBisCO and Prk were overexpressed, resulting in a strain that can incorporate CO_2_ into biomass using methanol as energy source, but no continuous growth on CO_2_ as sole carbon source was observed ^12^. So far, it was not possible to construct a synthetic autotrophic organism which can readily grow on CO_2_ as a sole carbon source to form its entire biomass.

The methylotrophic yeast *Pichia pastoris* (syn.: *Komagataella phaffii*) ^13,14^ is widely used as a production host for heterologous proteins serving both the biopharmaceutical ^15^ as well as the technical enzyme markets ^16^. It has gained increased interest as a chassis host for metabolic engineering ^17^ and as a model organism for studying peroxisome biogenesis ^18,19^. *P. pastoris* is capable of a methylotrophic lifestyle enabling it to use the C1 compound methanol as its sole energy and carbon source. After its oxidation to formaldehyde, methanol utilization is split into two branches: in the assimilatory branch methanol is fixated after oxidation to formaldehyde to finally yield phosphosugars needed for biomass generation, while in the dissimilatory pathway energy is produced in the form of NADH. Steps of the dissimilatory branch are carried out both in the peroxisome and in the cytosol, while the assimilatory branch is entirely localized in peroxisomes ^20^. Upon growth on methanol, peroxisomes are highly abundant within the cell and methanol utilization pathway associated enzymes are predominately expressed (e.g. alcohol oxidase 1 (Aox1)). Powerful genetic tools including CRISPR/Cas9 mediated techniques are available for *P. pastoris* ^17,21,22^ allowing to re-build and integrate entire metabolic pathways into this host without the use of integrative selection markers. In this work, we use *P. pastoris* as a chassis cell to *de novo* engineer a synthetic CBB cycle and to test its capability to promote autotrophic growth on CO_2_.

## 2. Results

### The xylulose-monophosphate (XuMP) cycle is used as a template to engineer a synthetic Calvin-Benson-Bassham (CBB) cycle into *P. pastoris*

The methanol assimilation pathway of *P. pastoris* is entirely localized in peroxisomes and shares high similarity with the CBB cycle ^20^. In both pathways a C1 molecule is transferred to a sugar phosphate generating a C-C bond. Methanol is oxidized by alcohol oxidase (Aox1 and Aox2 in *P. pastoris*) to formaldehyde. Subsequently, formaldehyde reacts with xylulose-5-phosphate (Xu5P) to dihydroxyacetone (DHA) and glyceraldehyde-3-phosphate (GAP) catalyzed by dihydroxyacetone synthase (Das1 and Das2 in *P. pastoris*). Xu5P is regenerated by pentose phosphate pathway (PPP) enzyme isoforms specifically enriched in peroxisomes upon cultivation on methanol ^20^. In total, three mole of methanol are converted to one mole of GAP, which can be used for biomass and energy generation. Likewise, in autotroph organisms, CO_2_ is added to ribulose-1,5-bisphosphate (RuBP) to form 3-phosphoglycerate (3-PGA) in a carboxylation reaction catalyzed by RuBisCO. 3-PGA is then phosphorylated and reduced to GAP. Remarkably, the XuMP cycle can be turned into a synthetic CBB cycle by addition of six enzymatic steps (i.e. all biochemical reactions of the CBB cycle present). The overall engineering strategy is visualized in Fig. 1. We considered a peroxisomal compartmentalization of the synthetic CBB cycle as beneficial, comparable to native C1 fixation in plant chloroplasts and in the XuMP cycle. To achieve this localization, a C-terminal peroxisomal targeting signal (PTS1) ^23^ was fused to each of the six engineered CBB enzymes. In a second control setup, all six heterologous genes were expressed in the cytosol without a PTS1 signal. The CBB cycle is split into 3 phases: carboxylation, reduction and regeneration ^24^. For the carboxylation reaction a RuBisCO form II from *Thiobacillus denitrificans* was integrated into the genome. The reduction phase to yield the triose phosphates, glyceraldehyde phosphate (*GAP*) and dihydroxyacetone phosphate (*DHAP*) was achieved by integration of the genes encoding for phosphoglycerate kinase (*PGK1*), glyceraldehyde-3-phosphate dehydrogenase (*TDH3*) from *Ogataea polymorpha* and triosephosphate isomerase (*TPI1*) from *Ogataea parapolymorpha*. The regeneration phase was completed by the addition of *TKL1* (encoding transketolase) from *O. parapolymorpha* and *PRK* (encoding phosphoribulokinase) from *Spinachia oleracea* to the five enzymes from the XuMP cycle (Fbp1, Fba1-2, Shb17, Rki1-2, Rpe1-2) already present in *P. pastoris*. For each molecule of CO_2_ fixated by this synthetic CBB cycle, 3 molecules of adenosine triphosphate (ATP) and 2 molecules of NADH are required. The energy is provided by methanol oxidation. The oxidation of one mole methanol yields two mole NADH. In order to separate the synthetic fixation machinery of CO_2_ from energy generation, we blocked the first steps of the methanol assimilatory pathway by deleting Das1 and Das2. To assist RuBisCO folding two molecular chaperones from *E. coli* (*GroEL* and *GroES*) were integrated into the genome ^8^. Furthermore, *AOX1* was deleted to reduce the formation rate of formaldehyde, which can still be formed by *AOX2* (encoding alcohol oxidase 2). In a nutshell, to enable a functional CO_2_ assimilation cycle in *P. pastoris*, three native genes were deleted and eight heterologous genes were integrated into the genome. Six of these genes are required for a fully functional CBB cycle and are either targeted to the peroxisome or to the cytosol. Two genes encode chaperones and are expressed in the cytosol. We generated these strains (listed in supplementary Table 1) in a three-step transformation procedure by a CRISPR/Cas9 mediated workflow for genome engineering in *P. pastoris* (see Fig. 2).

**Figure 1.**
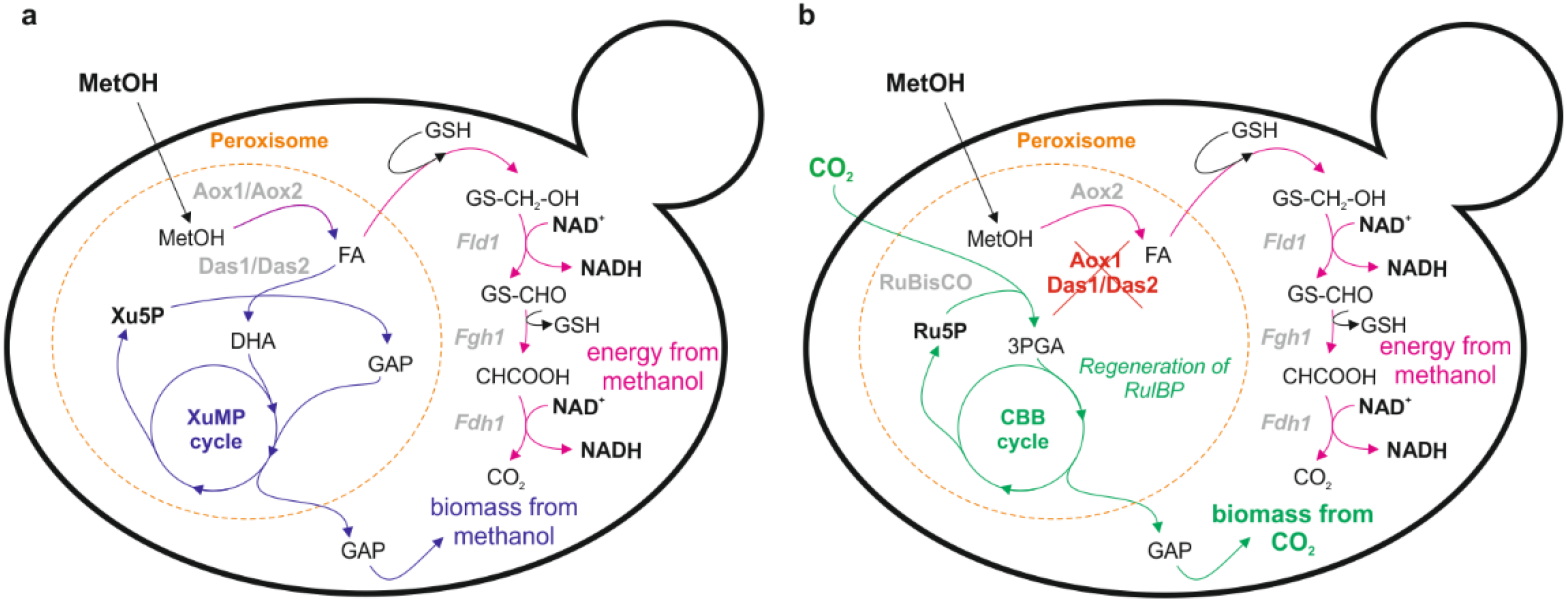
Scheme for engineering chemoorganoautotrophy in *Pichia pastoris*. (**a**) **Wild type *P. pastoris*** uses methanol (MetOH) for biomass generation in the assimilatory branch of the methanol utilization by fixation of formaldehyde (FA) to xylulose-5-phosphate (Xu5P) which is regenerated in the xylulose monophosphate (XuMP) cycle (purple pathway) and as an energy source in the dissimilatory branch (pink pathway) by oxidation to carbon dioxide (CO_2_) under formation of NADH. (**b**) In an **engineered strain** methanol assimilation is blocked by deletion of dihydroxyacetone synthase (*DAS1* and *DAS2*) and a synthetic CO_2_ fixation pathway similar to a Calvin-Benson-Bassham (CBB) cycle is integrated. RuBisCO carboxylates ribulose-1,5-bisphosphate (RuBP) which is regenerated in the synthetic CBB cycle; abbreviations: alcohol oxidase (Aox), formaldehyde dehydrogenase (Fld1), S-formylglutathione hydrolase (Fgh1), formate dehydrogenase (Fdh1), ribulose-1,5-bisphosphate carboxylase/oxygenase (RuBisCO), dihydroxyacetone (DHA), glyceraldehyde-3-phosphate (GAP), 3-phosphoglycerate (3PGA), glutathione (GSH), ribulose-1,5-bisphosphate (RuBP); A detailed depiction of the two pathways including all involved enzymes is given in Supplementary Figure 1.

**Figure 2.**
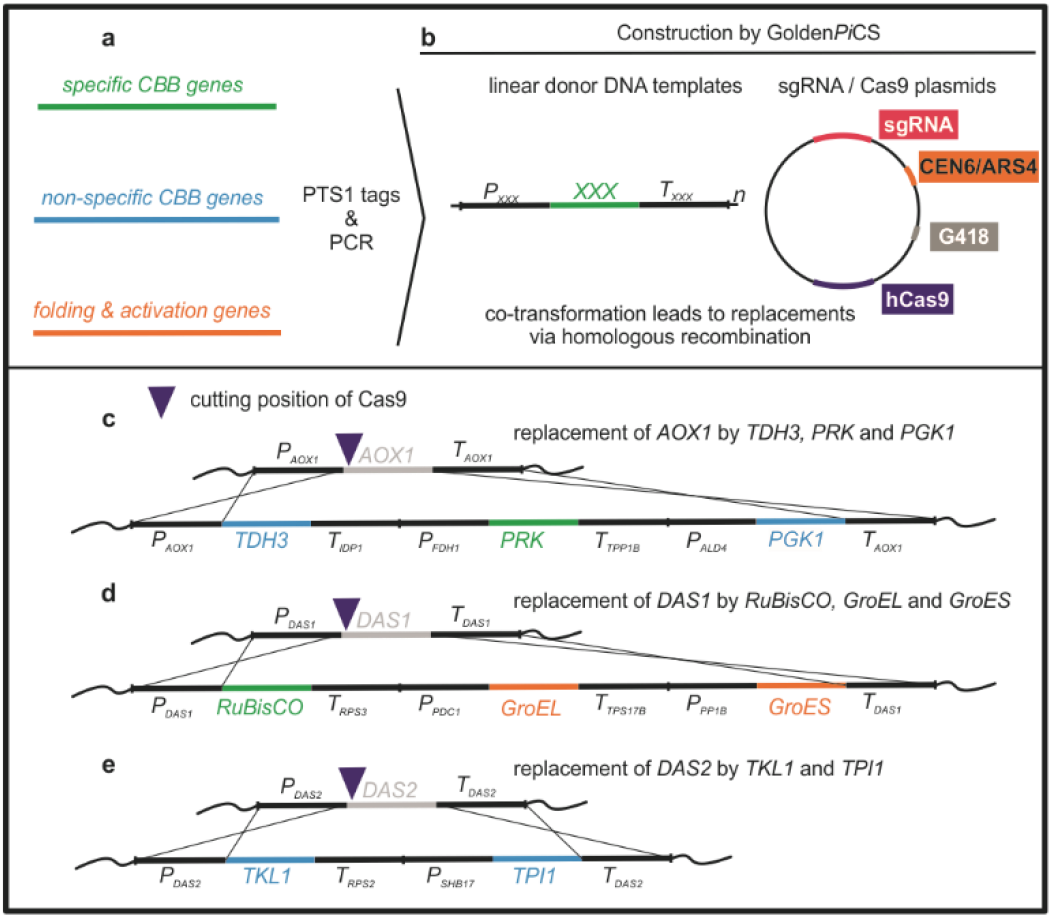
Engineering scheme for generation of *P. pastoris* strains with a synthetic CBB cycle. (**a-b**) **Plasmid construction**: (**a**) Genes for the proposed synthetic CBB cycle including CBB – specific (green – source are autotroph organisms), non-specific genes (blue – from *Ogataea* strains) and chaperones (from *Escherichia coli*) were amplified by PCR (*TDH3*, *PGK1*, *TPI1* and *TKL1*), or synthesized (*RuBisCO*, *PRK*, *GroEL* and *GroES*, codon optimized). The C-terminal PTS1 tag encoding for –SKL was added by PCR if needed. The entire gene names, source organisms and protein identifiers are summarized in Supplementary Table 4. (**b**) Generation of linear donor DNA templates and single guide RNA (sgRNA) / Cas9 plasmids; (**c** – **e**) **Homologous recombination (HR) mediated replacement of the native sequences** by co – transformation of a linear donor DNA with a sgRNA / Cas9 plasmid; replacement of (**c**) *AOX1* by an expression cassette for *TDH3*, *PRK* and *PGK1*, (**d**) *DAS1* by *RuBisCO*, *GroEL* and *GroES* and (**e**) *DAS2* by *TKL1* and *TPI1* was done consecutively in three transformation steps and is drawn schematically. The strains lacking one or more heterologous gene were constructed accordingly with altered donor DNA fragments. All strains constructed by this workflow are listed in Supplementary Table 1.

### Peroxisomal targeting of the CBB cycle leads to a higher growth rate on CO_2_ compared to a cytosolic localization

Strains expressing a cytosolic (CBBc+RuBisCO) and a peroxisomal (CBBp+RuBisCO) version of the CBB cycle were obtained. In order to test the ability of these strains to grow on CO_2_ as a carbon source, shake flask cultures were carried out in the presence of 5 % (v/v) CO_2_ (Fig. 3). A strain having the same genetic setup as the CBBp+RuBisCO strain but lacking RuBisCO (CBBpΔRuBisCO) served as a negative control. This strain showed no growth over the entire cultivation phase, whereas the CBBc+RubisCO and CBBp+RuBisCO strains were able to grow (Fig. 3). Furthermore, it was observed that the strains with a peroxisomal CBB pathway (CBBp+RuBisCO) grew significantly faster increasing from 0.35 ± 0.00 to 0.8 ± 0.08 g L^−1^ CDW in 164 hours compared to the strains with a cytosolic CBB cycle, which grew from 0.36 ± 0.02 to 0.5 ± 0.12 g L^−1^ CDW within the same timeframe.

**Figure 3.**
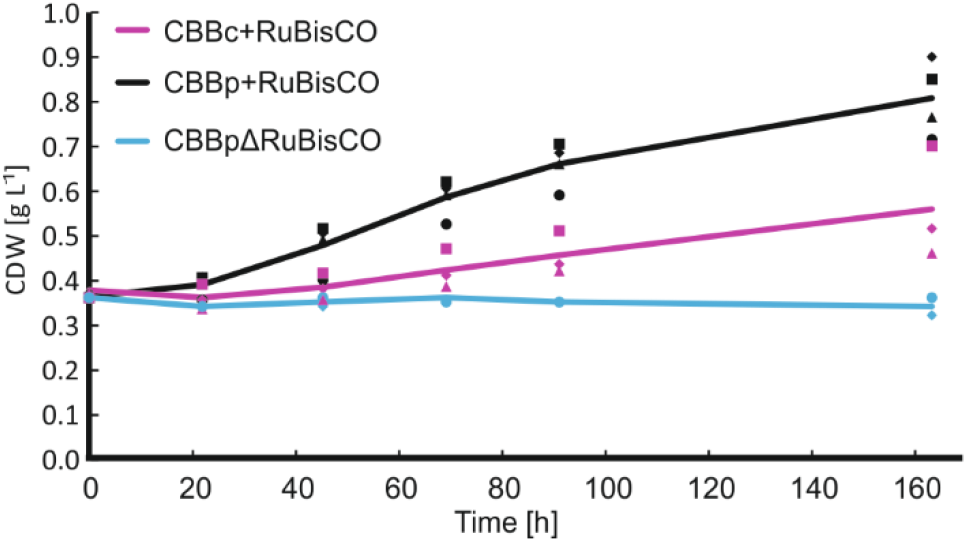
Peroxisomal targeting of the CBB pathway leads to increased growth. Cultivation of strains with cytosolic (CBBc+RuBisCO) and peroxisomal (CBBp+RuBisCO) pathway expression; Cell Dry Weight (CDW) values are calculated from OD_600_ measurements (correlation: 1 OD_600_ unit = 0.191 g L^−1^ CDW), for CBBpΔRuBisCO the mean of two technical replicates including single values is shown (blue line plus signs), for CBBp+RuBisCO (black line) and CBBc+RuBisCO (pink line) the means of at least three biological replicates (independent transformations) with single values (black and pink signs) is shown.

This experiment showed that strains expressing a complete CBB cycle can grow in the presence of CO_2_ and methanol, which indicates that the integrated CBB cycle is functional (Fig. 3). The superior growth behavior of the CBBp+RuBisCO strain confirmed that it is beneficial to compartmentalize the CBB cycle by targeting it to the peroxisome and thereby replacing the native XuMP cycle. As expected the negative control (CBBpΔRuBisCO) did not grow on media containing methanol and CO_2_. In this strain, the deletion of *DAS1* and *DAS2* prevents the methanol assimilation capability of *P. pastoris* and the RuBisCO is missing to complete a full CBB cycle. These results confirm that a deletion of *DAS1/DAS2* is sufficient to block the assimilation of methanol into biomass. The transketolase 1 (Tkl1) which is responsible for the conversion of GAP and S7P into R5P and Xu5P is a homolog of Das1 and Das2. Thus, Tkl1 might exhibit also a promiscuous activity for the reaction catalyzed by Das1/Das2, which is the conversion of formaldehyde and Xu5P to GAP and DHAP. However, since CBBpΔRuBisCO were not able to grow when supplemented with methanol, Tkl1 cannot substitute for Das1/Das2.

### *P. pastoris* strains expressing a functional CBB cycle in the peroxisome require CO_2_ for growth and biomass formation

In the following experiments the CO_2_ incorporation characteristics of the strains expressing a peroxisomal CBB cycle (CBBp+RubisCO) together with the negative control strain having a disrupted CBB cycle (CBBpΔRuBisCO) were tested in controlled bioreactor cultivations. After accumulation of biomass in a batch phase on glycerol, CBBp+RuBisCO and CBBpΔRuBisCO cells were fed with CO_2_ and/or methanol. Thereby, two different CO_2_ feeding regimes were used: in regime A, a set of both strains were first fed with methanol only and 0% (v/v) CO_2_ in the inlet gas and after 135 hours the CO_2_ was increased to 5 % (v/v). In regime B, a set of both strains was fed first with methanol and 5% CO_2_ and switched to 0% CO_2_ after 135 hours (Fig. 4). The bioreactors were operated at a high stirring- and gasflow-rate (1000 rpm and 35 sL h^−1^ respectively) to blow out CO_2_ formed by methanol oxidation. With these experimental settings, the CO_2_ concentration measured at the off-gas analyzer was below the detection limit of 0.1 % (v/v) also in cultures oxidizing methanol. Thus, a defined CO_2_ supply of the cells was controlled by changing the CO_2_ content of the inlet gas flow.

**Figure 4.**
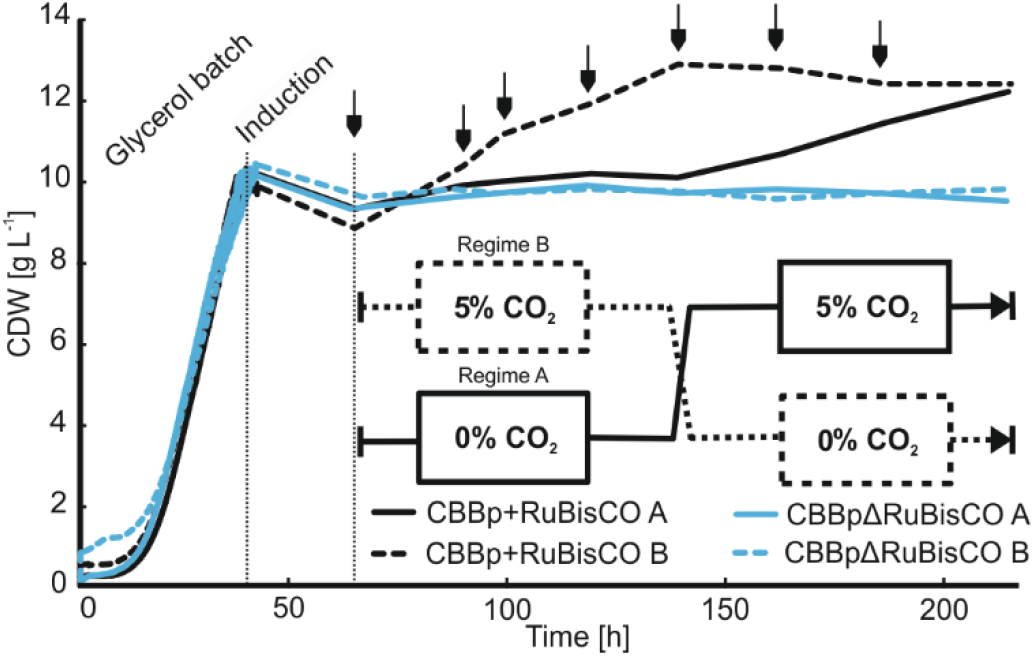
Growth in enginered CBBp+RuBisCO (A and B) depends on the supply of CO_2_ as a carbon source. Biomass formation in engineered CBBp+RuBisCO (solid (regime A) and dashed (regime B) black line) is shown compared to the control strain, which lack RuBisCO, GroEL and GroES (CBBpΔRuBisCO solid (A) and dashed (B) blue line. Cells were cultivated in batch phase on 16.0 g L^−1^ glycerol to a CDW of ~ 10 g L^−1^ and then induced with 0.5 % methanol (v/v) and 1 % CO_2_, afterwards methanol concentration was adjusted to 1 % (v/v) (arrows). After induction only CBBp+RuBisCO B and CBBpΔRuBisCO B were co-fed with 5 % CO_2_. After 3 days and occurrence of pronounced growth, the CO_2_ supply was set to 0 % for CBBp+RuBisCO B and CBBpΔRuBisCO B and increased to 5 % for CBBp+RuBisCO A and CBBpΔRuBisCO A. Cell Dry Weight (CDW) values were calculated from OD_600_ measurements (correlation: 1 OD_600_ unit = 0.191 g L^−1^ CDW).

The two strains (CBBp+RuBisCO and CBBpΔRuBisCO) growing under the CO_2_ feeding regime A (Fig. 4) did not grow without the addition of CO_2_. After commencing to feed with CO_2_ the strain CBBp+RuBisCO began to grow with a biomass formation rate of 0.029 g L^−1^ h^−1^ CDW. No growth was detected for the control strain CBBpΔRuBisCO. In regime B (Fig. 4), where CO_2_ is fed in the first phase, the CBBp+RuBisCO strain could grow with a rate of 0.036 g L^−1^ h^−1^ CDW and again no growth was detected for the negative control strain CBBpΔRuBisCO. In the second phase of this cultivation, where no CO_2_ was present in the inlet gas supply, no growth was detected for both strains. These results demonstrate that the CBBp+RuBisCO strains can take up and grow on CO_2_ as a carbon source and thus have a functional CO_2_ assimilation pathway. As energy source methanol can be oxidized by all strains (Supplementary Fig. 2). The control strains (CBBpΔRuBisCO) are lacking the essential step of the CBB cycle catalyzed by RuBisCO and thus the energy generated by methanol oxidation can only be used to sustain the energy requirements of cellular maintenance. In accordance, the methanol oxidation rate in CBBpΔRuBisCO strains with 0.027 g (MetOH) g (CDW)^−1^ h^−1^ was lower compared to CBBp+RuBisCO strains which have a methanol oxidation rate of 0.046 g (MetOH) g (CDW)^−1^ h^−1^ (Supplementary Fig. 2). The CBBp+RuBisCO strains only grew in the presence of both methanol and CO_2_. Even after a prolonged period on methanol as an energy carrier cells could start growing again, when CO_2_ is present in the inlet gas. Taken together these data demonstrate that the growth of the engineered strains expressing a synthetic CBB cycle in the peroxisomes of *P. pastoris* is dependent on the supply of CO_2_ as carbon source.

### Incorporation of CO_2_ into biomass is verified by ^13^C labelling

The dependency of CBBp+RuBisCO cells on the supply of gaseous CO_2_ for growth was shown in the previous experiment. However, to obtain further evidence that CO_2_ is taken up and incorporated into biomass, a study in which the labelling of biomass with ^13^C and ^12^C is measured by Elemental Analysis - Isotope Ratio Mass Spectrometry (EA-IRMS) was designed. The biomass of the engineered cells (CBBp+RuBisCO and CBBpΔRuBisCO) was enriched with ^13^C carbon by batch cultivation on minimal medium containing fully labelled ^13^C glycerol as the sole carbon source (Fig. 5 a). The ^13^C content in the biomass samples was measured by EA-IRMS. After the batch phase, the ^13^C enrichment reached 95 ± 0.5 % for the cultivation with CBBpΔRuBisCO and 97 ± 0.3 % for the CBBp+RuBisCO (Fig. 5 c). 100% ^13^C labelling cannot be reached, due to ^12^C carried over from the inoculum and the trace ^12^C contamination of the utilized ^13^C glycerol (^13^C enrichment >99 %). After the glycerol was consumed, the feeding regime was shifted to gaseous CO_2_ with a natural isotopologue distribution (98.901 ^12^C, 1.109 % ^13^C) and to methanol either with a natural isotopologue distribution or with ^13^C enrichment (>99 % ^13^C).

**Figure 5.**
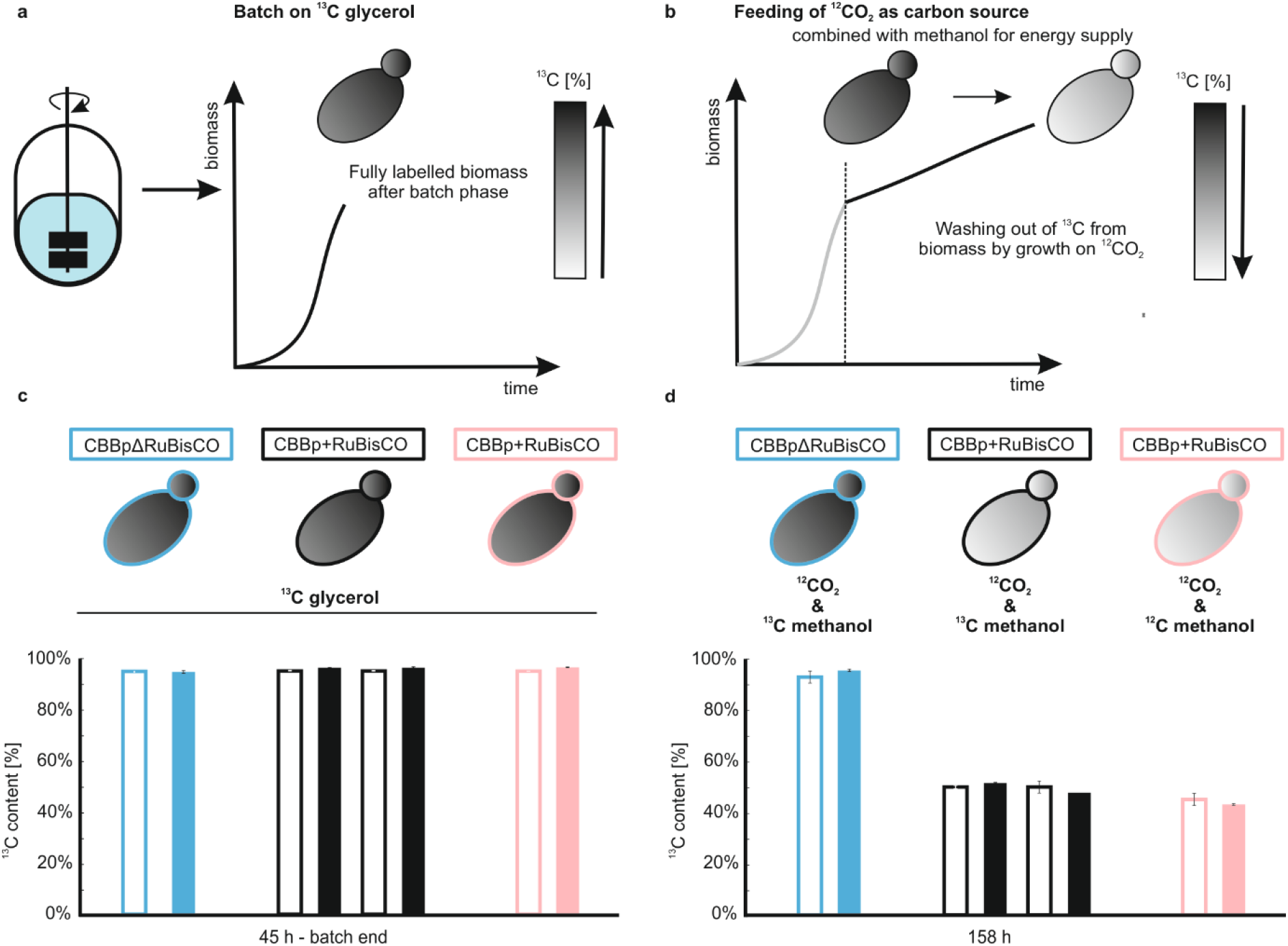
Labelling of engineered *P. pastoris* cells with ^13^C glycerol and ^12^C carbon dioxide. (**a**) CBBpΔRuBisCO and CBBp+RuBisCO (three technical replicates) cells were enriched for ^13^C by cultivation on fully labelled ^13^C glycerol as the sole carbon source to around 8.0 g L^−1^ CDW. White-to-black color schematically indicates the ^13^C [%] of the biomass; (**b**) After the batch phase the cells were fed with methanol (0.5 −1.0 % (v/v) and 5 % CO_2_ which led to growth in CBBp+RuBisCO and no growth for the CBBpΔRuBisCO. ^13^C methanol was used for two reactors containing CBBp+RuBisCO (black) and one reactor CBBpΔRuBisCO (blue), while ^12^C methanol was used for one reactor with CBBp+RuBisCO (pink). Incorporation of ^12^CO_2_ leads to a decrease of the total ^13^C content in the biomass. (**c** – **d**) Bar charts show total ^13^C content after the batch phase (45 h) (**c**) and after 158 h of cultivation (**d**), open bars show calculated values and filled bars show measured values obtained by EA-IRMS (error bars show *s.e.m* and for calculated values CDW *s.e.m.* was included).

As expected, CBBp+RuBisCO strains grew under the conditions with a mean biomass formation rate of 0.040 ± 0.003 g L^−1^ h^−1^ CDW, whereas CBBpΔRuBisCO strains showed no growth but still consumed methanol. During growth on CO_2_ and methanol the cells carried out approximately one cell doubling. Growth of the CBBp+RuBisCO strain was accompanied with a reduction of the ^13^C content in the biomass (Fig. 5 d). After 158 hours the ^13^C labelling content of the CBBp+RuBisCO fed with ^12^CO_2_ and ^13^C methanol reached 52 ± 0.3% or 48 ± 0.2% in a technical replicate. These data matched with the predicted values based on the measurement of accumulated biomass. The non-growing CBBpΔRuBisCO showed no change in the ^13^C labelling content over the entire cultivation period. In the fourth experiment, the CBBp+RuBisCO was cultivated with ^12^CO_2_ and ^12^C methanol. Also in this case the ^13^C labelling content decreased according to predicted values based on the biomass measurements. All values obtained in this experiment are shown in Supplementary Table 2.

These data show that the carbon coming from CO_2_ is incorporated into the biomass of a strain expressing a functional CBB cycle. The carbon from methanol is oxidized to CO_2_, but due to the high gas-flow rate, it is efficiently removed from the bioreactor. The negative control strain (CBBpΔRuBisCO), which can oxidize methanol but cannot incorporate CO_2_ into the biomass, showed consequently no change in the overall ^13^C labelling pattern. Taken together these data confirm that a *P. pastoris* strain containing a functional CBB cycle can utilize CO_2_ as a sole carbon source and methanol as energy source. *Per definitionem* the lifestyle of this strain can be described as chemoorganoautotrophic.

### Multiple cell doublings are possible by CBBp+RuBisCO strains using CO_2_ as the sole carbon source

In order to show that the chemoorganoautotrophic lifestyle of the engineered CBBp+RuBisCO can be maintained for multiple cell doublings, the strain was inoculated with a low cell density of 0.2 g L^−1^ CDW in a bioreactor and cultivated for 470 hours with a constant supply of methanol and CO_2_.

The growth profile is shown in Fig. 6. After induction, the cells quickly started to grow with an initial growth rate of µ = 0.020 ± 0.0014 h^−1^ in the first 20 h after inoculation. In the following the growth was reduced, but a continuous exponential growth phase between 100 up to 235 hours with a µ of 0.008 ± 0.0001 h^−1^ and a doubling time of 92 ± 1.7 h was observed. The higher initial growth rate could be attributed to the utilization of intracellular storage compounds, enzymes and metabolites which were accumulated during growth on the pre-medium.

**Figure 6.**
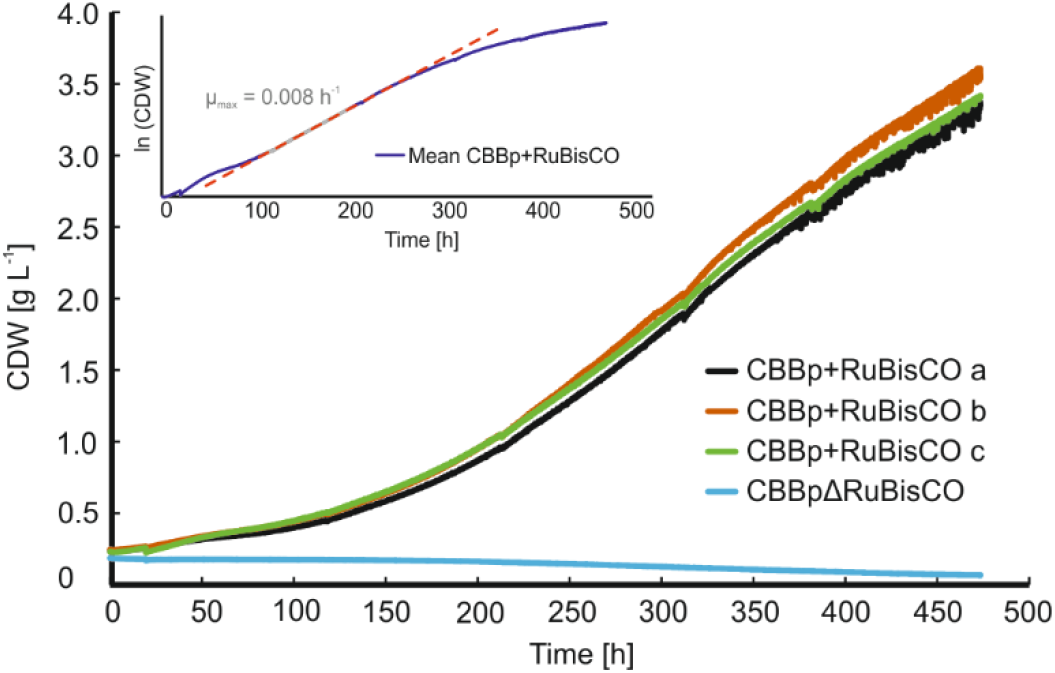
Bioreactor cultivation of CBBp+RuBisCO and CBBpΔRuBisCO strains inoculated at low cell density and grown in the presence of CO_2_. During cultivation, cells were fed with 5 % CO_2_ and methanol. Values for a single cultivation are shown for CBBpΔRuBisCO cells. For CBBp+RuBisCO cells (a - black, b - orange and c - green) CDW values of three biological replicates are shown. The insert shows the logarithmic mean CDW value of these three individual clones in purple (red dotted line shows linear fitting).

Therefore, the growth rate achieved in the prolonged exponential phase is determined as the µ_max_ of the culture under the given batch conditions. Over the entire observed 450 h of cultivation the cells performed a mean of 4.3 cell doublings. These data show that the engineered CBBp+RuBisCO strain can perform multiple cell doublings growing with a chemoorganoautotrophic lifestyle. The growth performance of the strains presented here is comparable to autotrophic organisms (Supplementary Table 3).

## 3. Conclusion

In this work, we reengineer yeast metabolism to enable a synthetic chemoorganoautotrophic lifestyle. Following a rational engineering strategy, a synthetic CBB cycle was introduced into the methylotrophic yeast *P. pastoris.* This enables the cell to utilize CO_2_ as a carbon source and to grow as an autotrophic organism. By splitting the metabolism into an energy generation module and a carbon fixation module, it is possible to decouple energy generation from carbon fixation ^6,12^. This modular design is crucial to demonstrate CO_2_ fixation by the engineered *P. pastoris* strains. In our design, methanol is used as an electron donor and is oxidized via formaldehyde to CO_2_ and NADH, which defines the lifestyle as chemoorganoautotrophic. Via the respiratory chain ATP is generated. The assimilatory pathway for methanol was blocked by deleting *DAS1* and *DAS2*. The obtained strain has the capability to perform multiple cell doublings on CO_2_ as sole carbon source. Our work demonstrates that the lifestyle of an organism can be switched from a chemoorganoheterotroph to a chemoorganoautotroph by metabolic engineering.

The synthetic CBB cycle is localized to the peroxisomes by targeting of the enzymes via PTS1 signals. This peroxisomal localization of the pathway enables superior growth on CO_2_ compared to a cytosolic localization. In contrast to *E. coli*, containing a cytosolic CBB pathway ^6^, no laboratory evolution was necessary to enable a functional CBB cycle. Engineering a CBB cycle into the yeast *S. cerevisiae* was shown to enable CO_2_ incorporation into ethanol but not into biomass ^8^. By supplementing the native methanol utilization pathway of *P. pastoris*, compartmentalization of the entire synthetic CBB cycle into the peroxisome was achieved, which can explain why CO_2_ fixation in *P. pastoris* can be achieved successfully, whereas it remains more challenging in other microorganisms. Thus *P. pastoris* is a promising chassis cell to incorporate and test CO_2_ fixation pathways *in vivo* including synthetic pathways like the malonyl-CoA-oxaloacetate-glyoxylate (MOG) pathway ^25^ or the crotonyl–coenzyme A (CoA)/ethylmalonyl-CoA/hydroxybutyryl-CoA (CETCH) cycle ^26^.

## 4. Materials and methods

### Plasmid construction

All expression cassettes and sgRNA / hCas9 plasmids were constructed by Golden Gate cloning ^22,27,28^. Three native genes of *P. pastoris* (*AOX1*, *DAS1* and *DAS2*) were replaced by the coding sequences of the genes listed in Supplementary Table 4. A CRISPR/Cas9 mediated homologous recombination (HR) directed system was used to construct the strains ^29^. Flanking regions needed for replacing the CDSs of *AOX1*, *DAS1* and *DAS2* were amplified from CBS7435 *wt* <u>g</u>enomic DNA (gDNA) (Promega, Wizard® Genomic DNA Purification Kit) by PCR (NEB, Q5® High-Fidelity DNA Polymerase). The other promoters (P_*ALD4*_, P_*FDH1*_, P_*SHB17*_, P_*PDC1*_ and P_*RPP1B*_) and terminators (T_*IDP1*_, T_*RPB1t*_, T_*RPS2t*_, T_*RPS3t*_ and T_*RPBS17Bt*_) were prepared accordingly and derived from genomic DNA of strain CBS7435 (Supplementary Table 5). CDSs of *TDH3* and *PGK1* were amplified from gDNA from *Ogataea polymporpha* (CBS 4732) and *TKL1* and *TPI1* from gDNA from *Ogataea parapolymorpha* (CBS 11895) according to the procedure described above and a C-terminal peroxisome targeting signal 1 (PTS1) ^23^encoding for SKL was added by amplification with primers carrying the additional nucleotides. The sequences encoding Prk (*Spinacia oleracea*), RuBisCO - cbbM (*Thiobacillus denitrificans*) and the chaperones GroEL and GroES (*E. coli*) were codon optimized (GeneArt, Regensburg, Germany). Final plasmids were digested with *BpiI* (*BbsI*), (ThermoFischer Scientific, USA) to obtain linear donor DNA templates used for replacement of the three native loci. Specific single guide RNAs (sgRNAs) were designed ^30^ and cloned into CRIS*Pi* plasmids ^29^. The genomic recognition sites for targeting the different loci with CRSIPR/Cas9 were CTAGGATATCAAACTCTTCG for *AOX1*, TGGAGAATAATCGAACAAAA for *DAS1* and CGACAAACTATAAGTAGATT for *DAS2.* The final sgRNA / Cas9 plasmids were checked by enzymatic digestion and by Sanger sequencing.

### Strain Construction

For cell engineering, *Komagataella phaffii (Pichia pastoris)* CBS7435 ^31,32^ was used as a host. Transformations were carried out as previously described ^15,29,33^. Primers listed in Supplementary Table 6 were used for detection of the correct replacement events of *AOX1*, *DAS1* and *DAS2* loci and correct integration was further verified by Sanger sequencing. All strains used in this study are listed in Supplementary Table 1.

### Cultivations in shake flask

Cultures were inoculated with a starting OD_600_ of 2.0 in YNB medium supplemented with 10 g L^−-1^ (NH_4_)_2_SO_4_. The flasks were incubated in a CO_2_ controlled incubator (5% CO_2_) shaking at 180 rpm at 30°C. The methanol concentration was adjusted up to 0.5 % (v/v) after the batch phase for induction and from there on maintained at 1% (v/v). Cell growth (OD_600_ and Cell Dry Weight (CDW) measurements) and metabolite profiles (HPLC analysis) were monitored ^34^.

### Bioreactor Cultivations

Single colonies were picked and used for inoculation of 100 mL pre-cultures in YPD medium (28°C o/n and 180 rpm). Bioreactors were inoculated with a starting OD_600_ of 1.0 or 0.19 g L^−1^ CDW. A defined medium consisting of citrate monohydrate (2 g L^−1^), glycerol (16 g L^−1^), MgSO_4_ 7H_2_O (0.5 g L^−1^), (NH_4_)_2_HPO_4_ (12.6 g L^−1^) KCL (0.9 g L^−1^) and biotin (0.0004 g L^−1^) was used. For the labelling experiment a medium containing H_3_PO_4_ (19.3 g L^−1^), CaCl_2_ 2H_2_O (0.025 g L^−1^), MgSO_4_ 7H_2_O (2.5 g L^−1^), KOH (2.0 g L^−1^), NaCl (0.22 g L^−1^), EDTA (0.6 g L^−1^), biotin (0.0004 g L^−1^) and fully labelled ^13^C glycerol (8.0 g L^−1^) was used. Both media were supplemented with trace elements. For the continuous growth fermentation YNB medium supplemented with 10 g L^−1^ (NH_4_)_2_SO_4_ was used.

The bioreactor cultivations were carried out in 1.4 L DASGIP reactors (Eppendorf, Germany). Cultivation temperature was controlled at 28 °C, pH at 5.0 by addition of 12.5 % ammonium hydroxide or with 5 M NaOH and 2 M HCl in case of YNB medium. The dissolved oxygen concentration was maintained above 20 % saturation. OD_600_ and cell dry weight (CDW) measurements as well as HPLC analysis to routinely measure glycerol, glucose, methanol and citrate concentrations ^34^ were carried out at every sample point. Induction was done by the addition of 0.5% methanol (v/v). The CO_2_ in the inlet gas was set to 1% during induction phase. After induction phase, the methanol concentration was adjusted to 1 - 1.5% (v/v) by bolus feeding and the CO_2_ concentration was set to 5 % throughout the cultivations. For the labelling experiment, the gas flow rate was set to 35 sL h^−1^ and the stirrer speed to 1000 rpm until the end of the fermentation.

### Elemental Analysis - Isotope Ratio Mass Spectrometry (EA-IRMS)

For the measurement of the biomass ^13^C content, the samples were taken during fermentation and processed at 4 °C. Volumes of cell suspension corresponding to approximately 0.5 mg of dried biomass was firstly washed with 0.1 M HCL and then twice with RO-H_2_O. Until analysis, the biomass samples were stored at −20 °C. The biomass material was homogenized and weighed in tin capsules. The ^13^C/^12^C ratio was measured with an Isotope Ratio Mass Spectrometer coupled to an Elemental Analyzer (EA-IRMS). EA-IRMS measurements were carried out by Imprint Analytics, Neutal, Austria.

## 5. Acknowledgements

This work has been supported by the Federal Ministry for Digital and Economic Affairs (bmwd), the Federal Ministry for Transport, Innovation and Technology (bmvit), the Styrian Business Promotion Agency SFG, the Standortagentur Tirol, Government of Lower Austria and ZIT - Technology Agency of the City of Vienna through the COMET-Funding Program managed by the Austrian Research Promotion Agency FFG. We also thank the Austrian Science Fund (FWF W1224—Doctoral Program on Biomolecular Technology of Proteins— BioToP). The funding agencies had no influence on the conduct of this research. EQ BOKU VIBT GmbH is acknowledged for providing fermentation equipment. We also kindly thank Hannes Rußmayer for help during labelling experiments.

## 6. Author Contributions

DM conceived and initiated the project. DM, MGS, TG, MS, and BG designed the experiments. TG carried out the experiments and together with MGS, MS and DM analyzed the data. TG, MGS and DM wrote the manuscript.

