## Supplemental Information for "A synthetic Calvin cycle enables autotrophic growth in yeast"

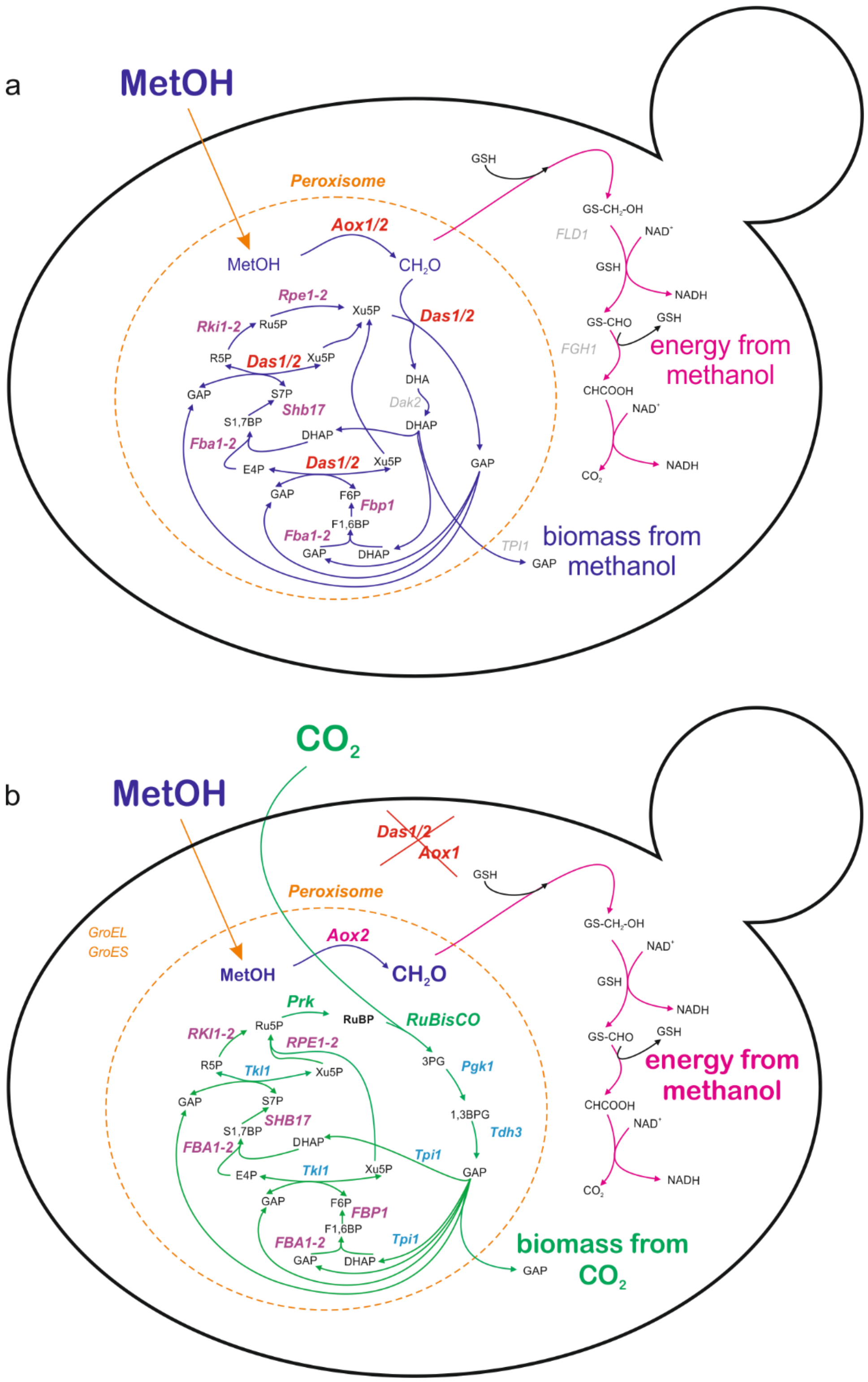


Supplementary Figure 1. **Description for engineering chemoorganoautotrophy in *Pichia pastoris*.** (**a**) **Wild type *P. pastoris*** is able to use methanol (MetOH) for biomass in the assimilatory branch of the methanol utilization by fixation of formaldehyde (FA) to xylulose-5-phosphate (Xu5P) which is regenerated in the xylulose monophosphate (XuMP) cycle (purple pathway) and as an energy source in the dissimilatory branch (pink pathway) by oxidation to carbon dioxide ($CO_2$) under formation of NADH. (**b**) In an **engineered strain** methanol assimilation is blocked by deletion of dihydroxyacetone synthase (*DAS1* and *DAS2*) and a $CO_2$ fixation pathway similar to a Calvin-Benson-Bassham (CBB) cycle is integrated (integration of *RuBisCO*, *PRK*, *TDH3*, *PGK1*, *TKL1*, *TPI1*, *GroEL* and *GroES*), RuBisCO carboxylates ribulose-1,5-bisphosphate (RuBP) which is regenerated in the synthetic CBB cycle; The enzymatic steps shown in magenta are present in both pathways (a and b); abbreviations: alcohol oxidase (Aox), dihydroxyacetone synthase (Das), fructose-1,6-bisphosphate aldolase (Fba1-2), fructose-1,6-bisphosphatase (Fbp1), sedoheptulose-1,7-bisphosphatase (Shb17), ribose-5-phosphate ketol-isomerase (Rki1-2), D-ribulose-5-phosphate 3-epimerase (Rpe1-2), formaldehyde dehydrogenase (Fld1), S-formylglutathione hydrolase (Fgh1), formate dehydrogenase (Fdh1), ribulose-1,5-bisphosphate carboxylase/oxygenase (RuBisCO), dihydroxyacetone (DHA), glyceraldehyde-3-phosphate (GAP), 3-phosphoglycerate (3PGA), glutathione (GSH), ribulose-1,5-bisphosphate (RuBP)

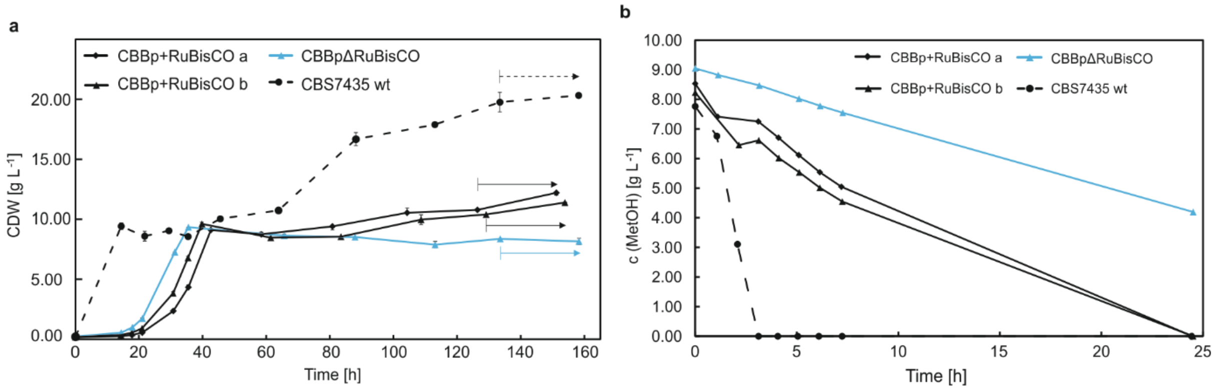


Supplementary Figure 2. (**a**) **Engineered CBBp+RuBisCO (a and b) are able to grow in presence of methanol and $CO_2$ while CBBpΔRuBisCO are not.** CBS7435 *wt* cells grow well in presence of both substrates, since methanol can be utilized for biomass and energy generation. Cells were cultivated in batch phase (16.0 g $L^{-1}$ glycerol) until 10 g $L^{-1}$ cell dry weight (CDW) and then fed with 0.5 – 1.0% (v/v) methanol pulses and a constant inflow of 5% $CO_2$. CDW values are calculated from OD measurements (correlation: 1 $OD_{600}$ unit = 0.191 g $L^{-1}$ CDW) and standard error bars indicate the standard error of 4 measurements; (**b**). **Methanol consumption.** Methanol uptake rates were determined during fermentation (timeframe of 24 h indicated by arrows in (a) and showed the highest methanol utilization by CBS7435 *wt* cells followed by the engineered CBBp+RuBisCO (a and b). The strain lacking RuBisCO (CBBpΔRuBisCO) showed slow methanol utilization.

*Supplementary Table 1 Strains used in this study*

| **Strain name** | **Genotype** |
|---|---|
| ***CBS 7435 Pichia pastoris*** | wt |
| **CBBp+RuBisCO** | Δ*aox1*::(*TDH3p, PRKp, PGK1p*)<br>Δ*das1::(RuBisCOp, GroELc, GroESc*)<br>Δ*das2*::(*TKL1p, TPI1p*) |
| **CBBpΔRuBisCO** | Δ*aox1*::(*TDH3p, PRKp, PGK1p*)<br>Δ*das1*<br>Δ*das2*::(*TKL1p, TPI1p*) |
| **CBBc** | Δ*aox1*::(*TDH3c, PRKc, PGK1c*)<br>Δ*das1::(RuBisCOc, GroELc, GroESc*)<br>Δ*das2*::(*TKL1c, TPI1c*) |

*Supplementary Table 2 Results from the labelling experiment*

| | **CBBpΔRuBisCO** | | **CBBp+RuBisCO I** | | **CBBp+RuBisCO II** | | **CBBp+RuBisCO III** | |
|---|---|---|---|---|---|---|---|---|
| **Strain** | **CBBpΔRuBisCO** | | **CBBp+RuBisCO** | | **CBBp+RuBisCO** | | **CBBp+RuBisCO** | |
| **C-source** | $^{12}CO_2$ / $^{13}CH_3OH$ | | $^{12}CO_2$ / $^{13}CH_3OH$ | | $^{12}CO_2$ / $^{13}CH_3OH$ | | $^{12}CO_2$ / $^{12}CH_3OH$ | |
| **Time [h]** | $^{13}C^{cal}$ | $^{13}C^{m}$ | $^{13}C^{cal}$ | $^{13}C^{m}$ | $^{13cal}$ | $^{13}C^{m}$ | $^{13cal}$ | $^{13}C^{m}$ |
| 45 Batch end | 96.3±0.1% | 95±0.5% | 96.5±0.2% | 97±0.1% | 96.5±0.2% | 97±0.3% | 96.4±0.0% | 97±0.1% |
| 85 | 96.3±1.5% | 95±0.8% | 80.3±0.3% | 76±0.9% | 76.3±3.0% | 77±0.6% | 68.8±2.2% | 72±0.2% |
| 133 | 96.3±0.1% | 95±0.3% | 57.4±0.6% | 57±0.6% | 56.8±2.6% | 58±0.1% | 52.8±1.9% | 50±0.4% |
| 158 | 96.3±2.3% | 96±0.5% | 50.3±0.6% | 52±0.3% | 50.0±2.4% | 48±0.2% | 45.4±2.3% | 43±0.4% |

*Supplementary Table 3 Comparison with enginnered and native autotrophic systems*

| **Strain** | **Specific growth rate μ [$h^{-1}$]** | **genotype** | **Energy source** | **Cultivation System** | **Reference** |
|---|---|---|---|---|---|
| *Methylobacterium extorquens AM1* | 0.003* | *engineered* | methanol | stirred reactor | [1] |
| *Eubacterium aggregans* | 0.005 | *wt* | hydrogen | syngas - anaerobic flasks | [2] |
| *Chlorella sp. NCTU-2* | 0.008 | *wt* | light | Bubble column | [3] |
| ***Pichia pastoris*** | **0.008** | ***engineered*** | **methanol** | **stirred reactor** | **This Study** |
| *Chlorella sp. NCTU-2* | 0.009 | *wt* | light | Centric tube | [3] |
| *Chlorella sp. NCTU-2* | 0.011 | *wt* | light | Porous centric tube | [3] |
| *Neochloris conjuncta* | 0.019 | *wt* | light | Bubble column | [4] |
| *Acetobacterium fimetarium* | 0.020 | *wt* | hydrogen | syngas - anaerobic flasks | [2] |
| *Neochloris texensis* | 0.022 | *wt* | light | Bubble column | [4] |
| *Neochloris terrestris* | 0.028 | *wt* | light | Bubble column | [4] |
| *Terrisporobacter mayombei* | 0.003 | *wt* | hydrogen | syngas - anaerobic flasks | [2] |
| *Botryococcus braunii* | 0.030 | *wt* | light | Bubble column | [4] |
| *Scenedesmus obliquus.* | 0.044 | *wt* | light | Bubble column | [4] |
| *Blautia hydrogenotrophica* | 0.066 | *wt* | hydrogen | syngas - anaerobic flasks | [2] |
| *Sporomusa acidovorans* | 0.071 | *wt* | hydrogen | syngas - anaerobic flasks | [2] |
| *Acetobacterium wieringae* | 0.081 | *wt* | hydrogen | syngas - anaerobic flasks | [2] |
| *Clostridium magnum* | 0.150 | *wt* | hydrogen | syngas - anaerobic flasks | [2] |
| *Sporomusa ovata* | 0.240 | *wt* | hydrogen | syngas - anaerobic flasks | [2] |

*limited to one cell doubling

*Supplementary Table 4 Genes used in this study including their sources*

| Gene Name | UniProt* | Source | EC Number | Full Name | PTS added |
|---|---|---|---|---|---|
| ***RuBisCOp/c*** | Q60028 | *Thiobacillus denitrificans* (ATCC 25259) | 4.1.1.39 | Ribulose-bisphosphate carboxylase | YES/NO* |
| ***PRKp/c*** | P09559.1 | *Spinacia oleracea* | 2.7.1.19 | Phosphoribulokinase | YES/NO* |
| ***PGK1p/c*** | A0A1B7SCV2 | *Ogataea polymporpha* (CBS 4732) | 2.7.2.3 | Phosphoglycerate kinase | YES/NO* |
| ***TDH3p/c*** | A0A1B7SCG5 | *Ogataea polymporpha* (CBS 4732) | 1.2.1.12 | Glyceraldehyde-3-phosphate dehydrogenase | YES/NO* |
| ***TPI1p/c*** | W1Q838 | *Ogataea parapolymorpha* (CBS11895) | 5.3.1.1 | Triosephosphate isomerase | YES/NO* |
| ***TKL1p/c*** | W1QKQ2 | *Ogataea parapolymorpha* (CBS11895) | 2.2.1.1 | Transketolase | YES/NO* |
| ***GroEL*** | B1XDP7 | *Escherichia coli (DH10B)* | N/A | molecular chaperone GroEL | NO |
| ***GroES*** | B1XDP6 | *Escherichia coli (DH10B)* | N/A | molecular chaperone GroES | NO |

*Supplementary Table 5 Promoters and terminators used in this study*

| Gene Name | Pxxx | Methanol Induced | location ID | Txxx | location ID | locus |
|---|---|---|---|---|---|---|
| ***PGK1p/c*** | $P_{ALD4}$ | Yes | PP7435_chr2 (1466285…1467148) | $T_{AOX1}$ | PP7435_chr4 (240891…241840 | *AOX1* |
| ***TDH3p/c*** | $P_{AOX1}$ | Yes | PP7435_chr4 (237941…238898) | $T_{IDP1}$ | PP7435_chr1 (1012481…1012975) | *AOX1* |
| ***TPI1p/c*** | $P_{SHB17}$ | Yes | PP7435_chr2 (340617…341606) | $T_{DAS2}$ | PP7435_chr3 (629173…630076) | *DAS2* |
| ***TKL1p/c*** | $P_{DAS2}$ | Yes | PP7435_chr3 (632201…633100) | $T_{RPS2}$ | PP7435_chr1 (2506918…2507385) | *DAS2* |
| ***RuBisCO p/c*** | $P_{DAS1}$ | Yes | PP7435_chr3 (634140…634688) | $T_{RPS3}$ | PP7435_chr1 (223093…223258) | *DAS1* |
| ***PRKp/c*** | $P_{FDH1}$ | Yes | PP7435_chr3 (423504…424503) | $T_{RPP1B}$ | PP7435_chr4 (463560…464058) | *AOX1* |
| ***GroEL*** | $P_{PDC1}$ | No | PP7435_chr3 (1860841…1861824) | $T_{RPS17B}$ | PP7435_chr2 (905111…905593) | *DAS1* |
| ***GroES*** | $P_{RPP1B}$ | No | PP7435_chr4 (462240…463233) | $T_{DAS1}$ | PP7435_chr3 (636813…637362) | *DAS1* |

*Supplementary Table 6: Primer sequences for verification of engineered loci, band sizes after PCR amplification are 4300 bp (AOX1), 3320 bp (DAS1) and 2420 (DAS2) for wt P. pastoris; Δaox1::TDH3, PRK, PGK1 (8700 bp); Δdas1::RuBisCO, GroEL, GroES (7200 bp), Δdas1 (3925 bp); Δdas2::TKL1, TPI1 (4700 bp)*

| **Primer** | **Sequence** |
|---|---|
| AOX1_Seq1_fw | TAATGTACATAGATCTTGAGATAAATTTCACGTTTA |
| AOX1_Seq1_rev | ATGAGACTGAGGTTTCATGAGTC |
| DAS1_Seq1_fw | ATTCTGTCGAAAATGGAAGCG |
| DAS1_Seq1_rev | CACTTGCATCACTGGCT |
| DAS2_Seq1_fw | CCTTTCCTTCCTGTTCCATT |
| DAS2_Seq1_rev | TCCTGAGTGACCGTTGTGTT |
